# Simulating the Dynamics of Targeted Capture Sequencing with CapSim

**DOI:** 10.1101/134510

**Authors:** Minh Duc Cao, Devika Ganesamoorthy, Lachlan J.M. Coin

## Abstract

**Motivation:** Targeted sequencing using capture probes has become increasingly popular in clinical applications due to its scalability and cost-effectiveness. The approach also allows for higher sequencing coverage of the targeted regions resulting in better analysis statistical power. However, because of the dynamics of the hybridisation process, it is difficult to evaluate the efficiency of the probe design prior to the experiments which are time consuming and costly.

**Results:** We developed CapSim, a software package for simulation of targeted sequencing. Given a genome sequence and a set of probes, CapSim simulates the fragmentation, the dynamics of probe hybridisation, and the sequencing of the captured fragments on Illumina and PacBio sequencing platforms. The simulated data can be used for evaluating the performance of the analysis pipeline, as well as the efficiency of the probe design. Parameters of the various stages in the sequencing process can also be evaluated in order to optimise the efficacy of the experiments.

**Availability:** CapSim is publicly available under BSD license at <u>https://github.com/mdcao/capsim</u>.

## Introduction

High-throughput sequencing (HTS) has tremendously revolutionised genomic studies for the ability for cost- and time-effective characterisation of the complete genetic information of a sample. In many clinical applications, only a panel of actionable regions are the subject for investigation [1, 4]. In these analyses, investigators often use a targeted capture sequencing protocol where a pool of synthesised oligonucleotides (probes) are used to selectively capture genomic fragments of interests using hybridisation. In an efficient design, only DNA fragments from the targeted loci are sequenced. This allows for deeper sequence coverage compared to whole genome sequencing at a much lower cost and faster time to results, resulting in a scalable approach for use in clinical laboratories.

Computational simulation has been indispensable in developing and benchmarking HTS data analysis tools [2]. Simulation data *in silico* are cheaper and faster to produce than real data; they are generated under controlled conditions and can be perfectly characterised. Furthermore, simulation also helps investigators to assess the performance of sequencing protocols, and to optimise the design prior to performing experiments. While numerous simulators are available for whole genome sequencing [2] and targeted exome sequencing [3], to the best of our knowledge, there is currently no existing tool, which can simulate the dynamics of the captured process *in silico*. We believe such a tool would be useful for assessing not only the computational analysis pipeline, but also the efficiency of a capture design.

Here, we present CapSim, a software package to meet the need for simulating targeted capture sequencing data. Given a set of probes, CapSim simulates the dynamics of probe hybridisation *in silico* to generate a set of fragments to be sequenced. Unlike most existing HTS simulation tools, CapSim emulates all various stages of the sequencing process, including fragmentation, fragment capture, and sequencing. Users can modify experimental parameters at each of these stages in order to optimise the sequencing protocols *in silico*. CapSim is written in Java and is able to run natively on any computing platform, making it easily accessible to bioinformatics community.

## Methods

The sequencing process starts with the fragmentation of the DNA, CapSim simulates this process by iteratively sampling fragments from the genome sequence with a given length distribution. We model the fragment length using a log-logistic distribution, which was found to provide the best fit to the fragment size distributions from several data sets (See Supplementary information). The parameters for the distribution are the median length (*α*) and the dispersion (*β*), making it intuitive for users to control the fragmentation simulation.

In the next step in targeted capture sequencing, the target probes bind to the fragments by hybridisation and the bound fragments are then pulled down by beads, which specifically binds to the DNA fragments hybridized to the capture probes. CapSim simulates this process by mapping the probes to the fragments. To be computationally efficient, CapSim first maps the probes to the genome sequence and once a fragment is sampled from the genome, it uses a greedy algorithm to determines the maximum number of probes that can bind to the fragment. We use a random variable to model the stochastic nature of the capture process in which the probability of a fragment being captured is proportional to the number of probes bound, and is inversely proportional to the length of the fragment. This simulation of the dynamics of the hybridisation is shown to generate more realistic captured sequencing data than Wessim [3], the only existing tool for simulating captured sequencing (See Results).

The captured fragments are then subjected to *in silico* sequencing. For Illumina sequencing, the adapter ligated to the DNA fragments binds to the oligonucleotides attached to the flow-cell, thereby forming clusters of fragments in the flow-cell. As the clustering introduces biases toward certain fragment size (see Supplementary information), CapSim introduces a size distribution of fragments that form clusters for sequencing. It uses a Log-Logistic distribution to sample from fragments simulated from the capture step. These selected fragments are then used for sequencing simulation in which reads are copied from the two ends of the fragments with errors introduced.

For PacBio sequencing, two hairpin adaptors are ligated to both ends of a double-stranded DNA fragment, creating a single-stranded closed loop, called SMRTBell. In sequencing the fragment, the SMRTBell traverses through a polymerase enzyme attached to the bottom where incorporated nucleotides are detected via fluorescence. If the polymerase read (the whole read from the well) is longer than the fragment, both strands of the fragment are sequenced in multiple passes. CapSim simulates this process by sampling polymerase read length from a given distribution. In simulating the sequencing of a fragment, CapSim alternates copying the two strands with a PacBio error profile, until reaching the polymerase read length.

We used Agilent SureSelect Target Enrichment kit to enrich the targeted regions and performed capture sequencing on NA12878 sample on both Illumina and PacBio sequencing platforms (See supplementary Information).

## Results

Fig. 1 shows the comparison in sequence read distribution between simulation data and real capture sequencing data on NA12878 sample on both Illumina and PacBio platforms, in a region of chromosome 17 containing the ARBB2 gene. Read distribution from CapSim simulation data closely resembles the real sequencing data from both platforms. Especially, the simulation data for both platforms show a region of off-target capture (1kb upstream of the target region) which was detected on the real sequencing data. In comparison simulation data from Wessim [3], the only existing captured sequencing simulator, did not replicate the sequence read distribution of real capture sequencing data and the off-target capture was not detected.

**Figure 1:**
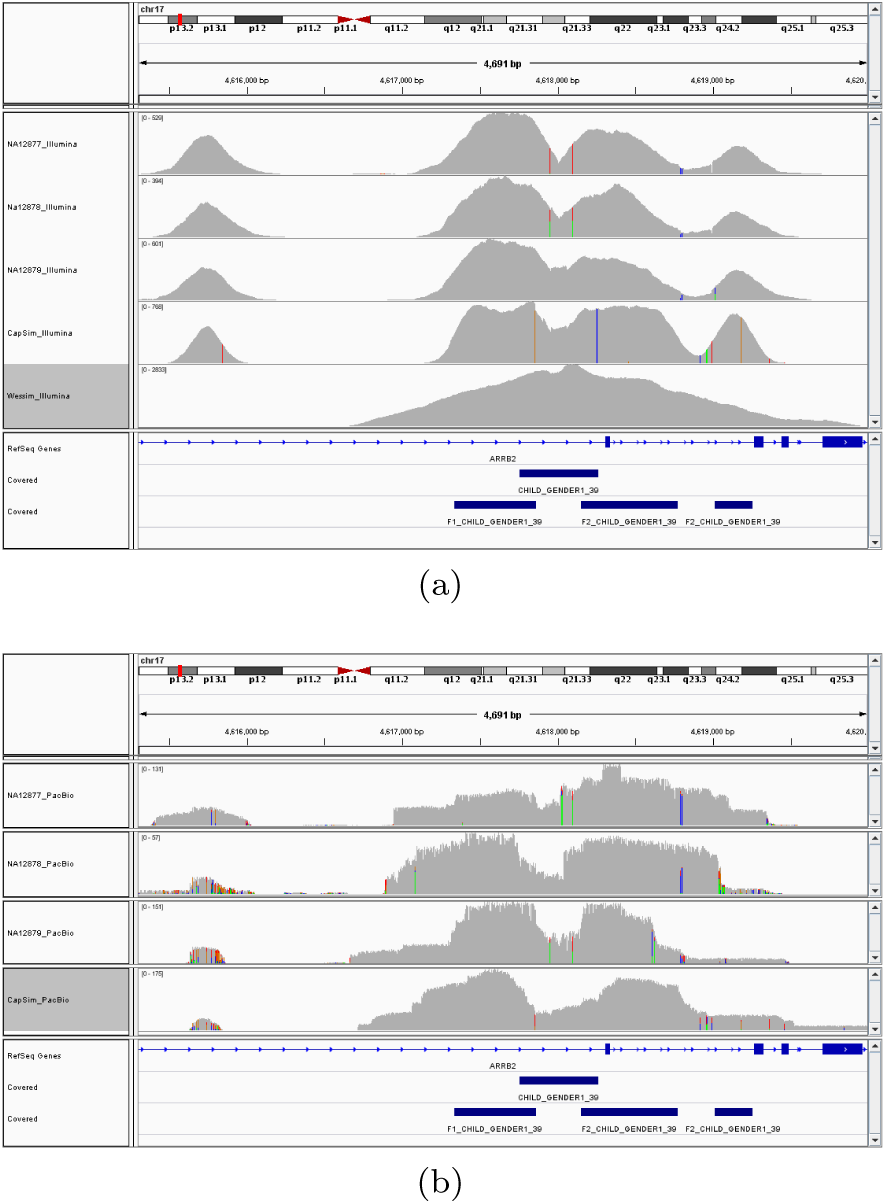
Depth coverage from capture simulation data compared to real sequencing data from Illumina MiSeq (a) and PacBio (b) platforms. Real data are shown in the top three panels and simulation data from CapSim are in the fourth panel. Simulation data from Wessim are shown in fifth panel for Illumina sequencing. Position of the capture probes are shown in blue in the bottom capture track panels.

## Conclusion

In this manuscript we have introduced CapSim a tool for simulation of targeted capture sequencing on both Illumina and PacBio platforms. Unlike most existing HTS data simulators, CapSim simulates the generation of data at each intermediate stage of the capture sequencing process. It provides parameters for controlling these stages, allowing users to evaluate the effects of the experiment design before running the experiment. Furthermore, CapSim will allow users to assess the efficiency of the probes for on-target capture regions and will reduce the off-target capture.

